# Tumor intrinsic regulation of PD-L1 and of interferon Type I *via* an SLC25A1-driven mitochondrial pathway, influences the anti-tumor immune response

**DOI:** 10.1101/2025.09.20.677512

**Authors:** Rami Mosaoa, Maha Moussa, Aditya Kavuturu, Salan Preet Kaur, Garrett Graham, Cecil Han, Chris Albanese, Marta Catalfamo, Maria Laura Avantaggiati

## Abstract

Immune checkpoint inhibitors (ICIs) have transformed cancer therapy, but variable patient responses highlight the need to better regulators of immune sensitivity. Here, we identify the mitochondrial citrate carrier SLC25A1 as a determinant of anti-PD-L1 antibody therapy responsiveness through a dual regulation of type I interferon (IFN-I) signaling and of PD-L1 expression. SLC25A1 promotes a mitochondrial-to-nuclear retrograde signaling *via* cytosolic accumulation of mitochondrial DNA, activation of the cGAS-STAT1 axis, and establishment of a virus mimicry state that triggers the IFN-I response. This activation is enriched in cancer stem cell populations, consistent with the role for SLC25A1 in tumor stemness and therapy resistance. Moreover, SLC25A1 also regulates PD-L1 protein levels through a newly identified fumarate-Keap1-PD-L1 axis, whereby fumarate inhibits Keap1, leading to PD-L1 up-regulation. In vivo, tumors expressing high levels of SLC25A1 exhibit an inflammatory environment and increased sensitivity to PD-L1 blockade, but accelerated growth in the absence of anti-PD-L1 treatment. These findings position SLC25A1 as a novel regulator of mitochondrial-driven IFN-I signaling and PD-L1 stability, and suggest that SLC25A1 exploits PD-L1 to evade immune surveillance, while at the same time creating an intrinsic tumor vulnerability to checkpoint blockade. Thus, SLC25A1 may serve both as a biomarker of response and as a target to enhance the efficacy of immunotherapy.

## INTRODUCTION

Immune checkpoint inhibitors (ICIs) have reshaped therapeutic strategies for various malignancies, particularly lung cancer and melanoma [1–3]. Although durable and objective responses are observed in a subset of patients, a considerable number of patients do not respond to treatment, limiting the overall clinical benefit. Several factors influence the response to ICIs, including the tumor immunogenicity and the composition of the tumor immune microenvironment (TIME). Favorable responses are often associated with an inflamed TIME, characterized by infiltration of CD8⁺ T cells, dendritic cells, and M1 macrophages, along with expression of Major Histocompatibility Complex (MHC) class I and II molecules. Conversely, “cold” tumors, which exhibit minimal immune cell infiltration and reduced antigenicity, are less responsive to ICIs [4–6]. Consequently, strategies aimed at converting cold tumors into inflamed tumors are a focus of ongoing investigations.

Given that tumor cells frequently up-regulate PD-L1 as a mechanism to evade immune surveillance, understanding the regulation of PD-L1 stability is also of significant interest. High PD-L1 expression often correlates with improved responses to ICIs, and inhibition of PD-L1 degradation pathways in certain tumor contexts has been proposed as a strategy to enhance immunotherapy efficacy. For instance, CDK4/6 inhibitors (e.g., palbociclib) have been shown to stabilize PD-L1 and to synergize with PD-L1 blockade [7–10]. Conversely, in tumors with sufficient immunogenicity, promoting PD-L1 degradation may facilitate the recruitment of effector immune cells and enhance tumor rejection [11]. Thus, elucidating the mechanisms regulating PD-L1 turnover has clinical relevance.

Type I interferons (IFN-I), primarily represented by IFNα and IFNβ, are key mediators of the immune response to infections and also, have complex roles in cancer biology [12–16]. During viral infections, cytosolic viral nucleic acids activate IFN-I signaling, leading to the transcriptional induction of interferon-stimulated genes (ISGs) that mediate antiviral and inflammatory responses. In cancer, the effects of IFN-I are context-dependent. Certain ISGs, classified as the DNA damage resistance signature (IRDS), are over-expressed in various tumor types and have been implicated in promoting cancer stemness, radiation resistance, and resistance to apoptosis mediated by CD95 (Fas) [17–20]. Similarly, in the context of ICI therapy, IFN-I signaling exhibits dual roles. On one hand, sustained IFN-I activation has been associated with therapeutic resistance, partly through feedback loops that induce PD-L1 expression and contribute to T cell exhaustion [20,21]. This regulatory mechanism, which serves to limit immune-mediated tissue damage during infections, can be co-opted by tumors to promote immune evasion. On the other hand, IFN-I signaling has also been reported to enhance anti-tumor immunity, and systemic administration of IFNα and IFNβ has been shown to improve responses to immunotherapy in melanoma and other tumors [21–24]. Significantly, proteomic analyses in melanoma patients showed that the IFN-I pathway activation correlates with favorable response to ICI treatment, owing to mitochondrial oxidative phosphorylation signatures, but the underlying mechanisms remain incompletely defined [25]. In the same study, up-regulation of the mitochondrial citrate carrier, SLC25A1, appeared to correlate with the improved response to immunotherapy.

Recent evidence suggests that cancer cells can autonomously activate IFN-I pathways through a “virus mimicry” state, induced by the aberrant presence of endogenous nucleic acids in the cytosol [26,27]. A virus mimicry state can be therapeutically triggered by DNA demethylating agents, which reactivate normally silenced transposable elements, selectively affecting cancer stem cell populations [27,28]. Similarly, radiation therapy induces IFN-I responses through DNA damage-mediated cytosolic DNA accumulation, potentially enhancing the efficacy of immunotherapy [29]. Low levels of cytosolic DNA appear to be a common feature in many cancers, leading to chronic, low-level IFN-I signaling [30]. Furthermore, the cytoplasmic presence of micronuclei resulting from chromosomal segregation errors can activate IFN-I pathways and has been associated with increased metastatic potential [31]. Collectively, these findings suggest that the context, the upstream triggering signals, as well as the magnitude and quality of IFN-I activation are critical determinants of its effects on tumor progression and therapeutic outcomes. However, it remains incompletely clear which tumor-intrinsic factors, independent of exogenous therapeutic interventions, are responsible for inducing virus mimicry states and regulate PD-L1 expression.

We have shown before that the main function of SLC25A1 consists of transporting citrate bi-directionally in and out of the mitochondria, while bringing malate inside [32–35]. Our previous mechanistic studies, which also involved the development of the most potent SLC25A1 inhibitor, CTPI-2, demonstrated that by regulating the flux of citrate/malate across the mitochondria, SLC25A1 promotes OXPHOS, maintains adequate TCA cycle flux, and allows metabolic adaptation promoting tumor progression [33,34]. We now show here that SLC25A1 regulates the response to anti-PD-L1 therapy and recruits IFN-I signaling. We also uncover a novel fumarate–Keap1–PD-L1 regulatory axis and propose that tumors with high levels of expression of SLC25A1 and IFN-I signaling have intrinsic vulnerability to checkpoint blockade.

## RESULTS

### SLC25A1 elicits a mitochondrial-to-nucleus retrograde signaling that induces the antiviral IFN-I response

We have shown before that SLC25A1 is frequently, but not universally expressed in several cancer cell lines and in metastatic lymph node biopsies derived from patients affected by Non-Small Cell Lung Cancer (NSCLC) [34]. To expand these observations, we interrogated a tumor array platform derived from NSCLC and patient-derived cancer cells [34]. As shown in Fig. 1A, the expression of SLC25A1 is variable in NSCLC, with some tumors showing elevated levels and others with intermediate or no expression. Similarly, SLC25A1 expression was elevated in some patient-derived tumors relative to normal adjacent tissues, but was completely absent in a subset of the patient-derived cells (e.g., T3, T6 and T7; Fig. 1 B).

**Fig. 1:**
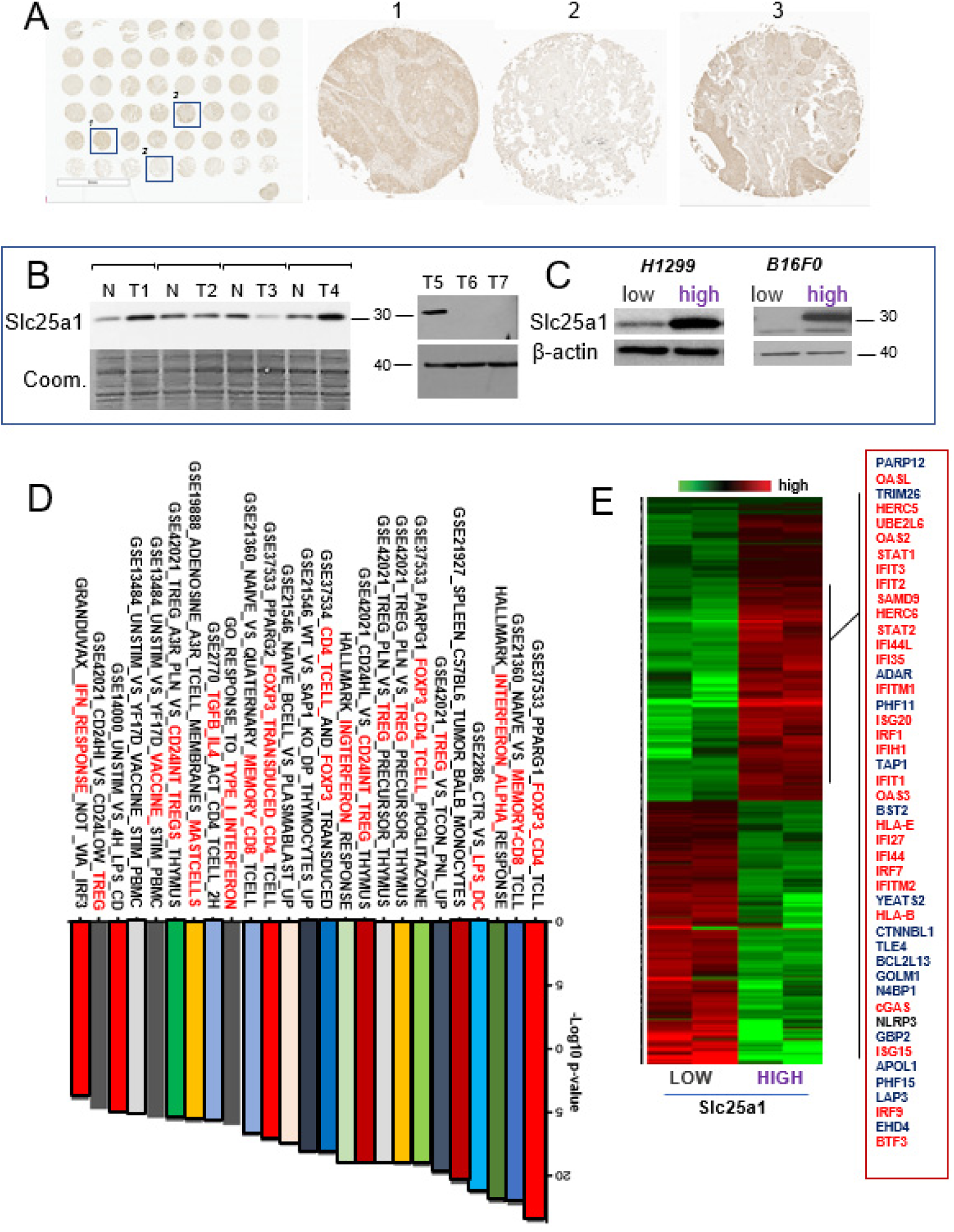
An immune-related signature is activated in SLC25A1-expressing NSCLC and melanoma spheroids. **A.** Immuno-histochemistry (IHC) of SLC25A1 in NSCLCs derived tumors. Squares indicate enlarged tumors shown in panels 1, 2 and 3. **B.** SLC25A1 levels in normal (N) tumor (T) tissues derived from patients with NSCLCs. **C.** Protein expression levels of SLC25A1 in H1299 and B16F0 cells naive (low), or expressing exogenous SLC25A1 (high). **D.** Gene set enrichment analysis (GSEA) of microarray experiments revealed significant enrichment of immune- and metabolism-related pathways in SLC25A1^high^ versus H1299-SLC25A1^low^ spheroids. Bars represent enriched gene sets ranked by statistical significance (–log10 *p*-value). Pathways associated with interferon signaling, immune checkpoint regulation, and T cell responses are highlighted in red. **E.** Heatmap of most enriched genes identified by Illumina arrays in H1299 control or SLC25A1-expressing spheres representing the ISGs. In red are the genes validated in this study with qRT-PCR or immunoblot.

To identify the pathways that are directly under the control of SLC25A1, we initially employed H1299 lung cancer or B16F0 melanoma cell lines (both of which have very low SLC25A1 expression), which were stably transfected with a vector encoding for human SLC25A1, resulting in moderately high expression of the protein (SLC25A1^high^), Fig. 1C. To identify the transcriptional program driven by SLC25A1, we then performed transcriptomic profiling in SLC25A1^low^ or SLC25A1^high^ H1299 cells grown in spheroid conditions, to model the three-dimensional architecture of tumors. Notably, the results of the gene expression arrays revealed induction of many genes and signal pathways involved in the regulation of the anti-tumor immune response. These included a Foxp3 signature that controls the development and function of regulatory T (Treg) cells, Treg signal pathways and others (Fig. 1D). Among these signatures, the IFN-Iα and IFN-I response were very robustly induced and included a broad spectrum of Interferon-Stimulated Genes (ISGs), as well as the cytosolic DNA sensors GMP-AMP synthase, cGAS (Fig. 1E). The major histocompatibility complex class I (MHC-I) *HLA-A*, *HLA*-*B*, and *HLA*-*E* were also up-regulated, likely due to IFN-I activation and/or enhanced immunogenicity of SLC25A1-enriched spheroids.

We then validated the expression array data by assessing the mRNA levels of members of the ISG family with qRT-PCR. This approach confirmed the induction of a large gene set typical of the IFN-I response in SLC25A1-expressing spheroids (Fig. 2A). To rule out that the induction of the IFN-I response is a cell type-specific phenomenon, we assessed the ISGs mRNA levels in murine melanoma B16F0 SLC25A1^high^ cells. Similar to H1299 cells, we observed induction of the ISGs by SLC25A1 in B16F0 as well (Fig. 2B). Similar results were also obtained in the kidney embryonic carcinoma 293T cell line (not shown). Further, the previously identified SLC25A1 inhibitor, CTPI-2, blunted the expression of ISGs in naïve H1299 spheres (Supplementary Fig.1A). In addition, Mouse Embryonic Fibroblasts (MEFs) lacking one *Slc25a1* allele (*Slc25a1+/-*) display reduced expression of ISGs relative to wild type MEFs (Supplementary Fig. 1B). In addition, as expected, both cGAS and STAT1 protein levels were strongly induced by SLC25A1 (Fig. 2C).

**Figure 2.**
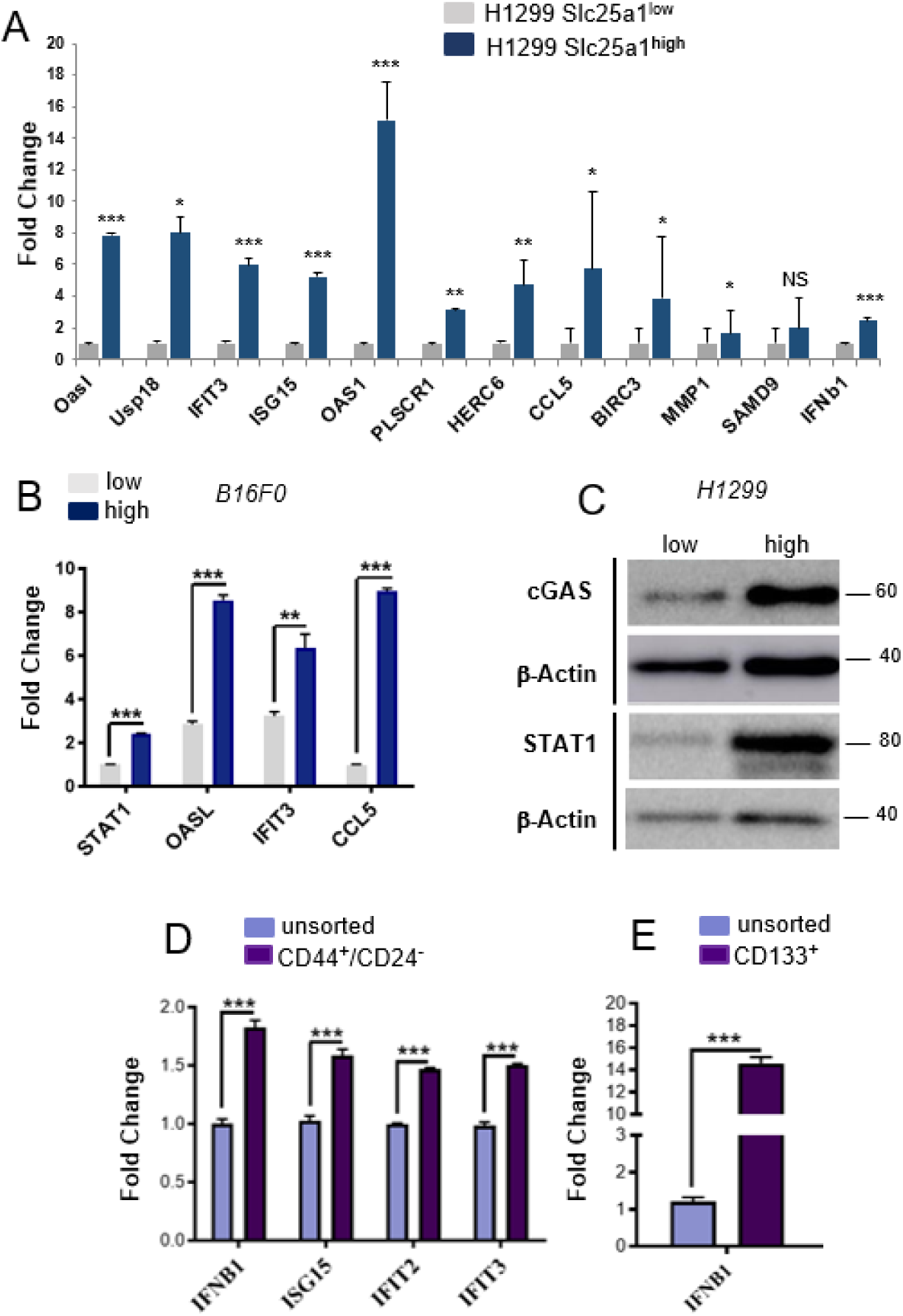
SLC25A1 conveys an IFN-I antiviral response. **A.** Fold change in gene expression of the indicated ISGs in the indicated H1299 spheres. **B.** Fold change in gene expression of the indicated ISGs in B16F0 spheres **C.** Levels of c-GAS, STAT1 and -Actin in SLC25A1 H1299 low and high cells. **D-E**. Fold change in gene expression of the indicated ISGs in CD44^+^/CD24^-^ CSCs (D), or CD133^+^ (E) CSCs that were isolated from SLC25A1^high^ spheres, relative to control negative populations. The error bars represent standard deviations, asterisks refer to *\*p* ≤ 0.05; *\*\*p* ≤ 0.01*; ***p* ≤ 0.001 by unpaired *T*-test.

Given recent evidence suggesting that activation of IFN-I signaling particularly in the cancer stem cell population (CSC), and our previous work demonstrating that SLC25A1 favors the expansion of the CSCs [17,34], we next explored whether SLC25A1 drives activation of the IFN-I pathway specifically within this population. Similar to embryonic and adult stem cells, CSCs from different cancer types can be identified and isolated based on the expression of stemness-associated surface markers, including CD133, CD44, and low CD24. To test the hypothesis that IFN-I activation is preferentially enriched within CSCs, SLC25A1^high^ spheroids were cultured for three days, followed by isolation of CSCs using fluorescence-activated cell sorting (FACS) based on surface expression of CD133, CD44, and CD24. The expression of interferon-stimulated genes (ISGs) in the isolated CSC population was subsequently analyzed by qRT-PCR. The results of these experiments demonstrated that IFNb1 and a subset of ISGs were more prominently expressed in the CD133-positive cell population compared to CD133-negative cells as well as in the CD44-positive/CD24-negative population (Fig. 2 D,E).

These findings reveal, for the first time, a physiological role for SLC25A1 in regulating IFN-I signaling in both normal cells and tumor CSCs. They further identify this protein as a key mediator of mitochondria-to-nucleus retrograde signaling, an essential communication pathway through which mitochondria transmit signals to the nucleus [36,37].

### Cytosolic mtDNA accumulation by SLC25A1 is responsible for IFN-I activation and induction of a virus mimicry state

The IFN-I response is activated under normal physiological conditions by the presence of exogenous (e.g., viral) or endogenous nucleic acids in the cytosol, which are in turn sensed by the Pattern Recognition Receptors (PRRs), including cGAS, which specifically recognizes cytosolic DNA [26,38,39]. This activation leads to a cascade of events involving the induction of Interferon Regulatory Factor 3 and 7 (IRF3/7), of interferon-induced protein with tetratricopeptide repeats 2 (IFIT2), which culminates in the induction of IFN-⍺ and IFN-β. These cytokines initiate autocrine and paracrine signaling loops by binding to their receptors (IFNAR1/2) and activating the Janus Kinase/Signal Transducers and Activators of Transcription (JAK/STAT) pathway, ultimately resulting in the amplification of type I interferon (IFN-I) signaling *via* STAT proteins. We therefore sought to investigate whether SLC25A1 induces this pathway by promoting mtDNA accumulation in the cytosol. To this end, we used primers designed to amplify regions of mitochondrial genes including *Cytochrome B (CYTB), NADH Dehydrogenase (ND2) and 16S ribosomal RNA gene (16S)*. The scheme of the mtDNA is shown in Supplementary Fig. 2A and the position of the primers employed is shown therein (closed balls). We found a significant increase in the levels of cytosolic mtDNA in both H1299 and 293T SLC25A1^high^ cells (Fig. 3A,B).

**Figure 3.**
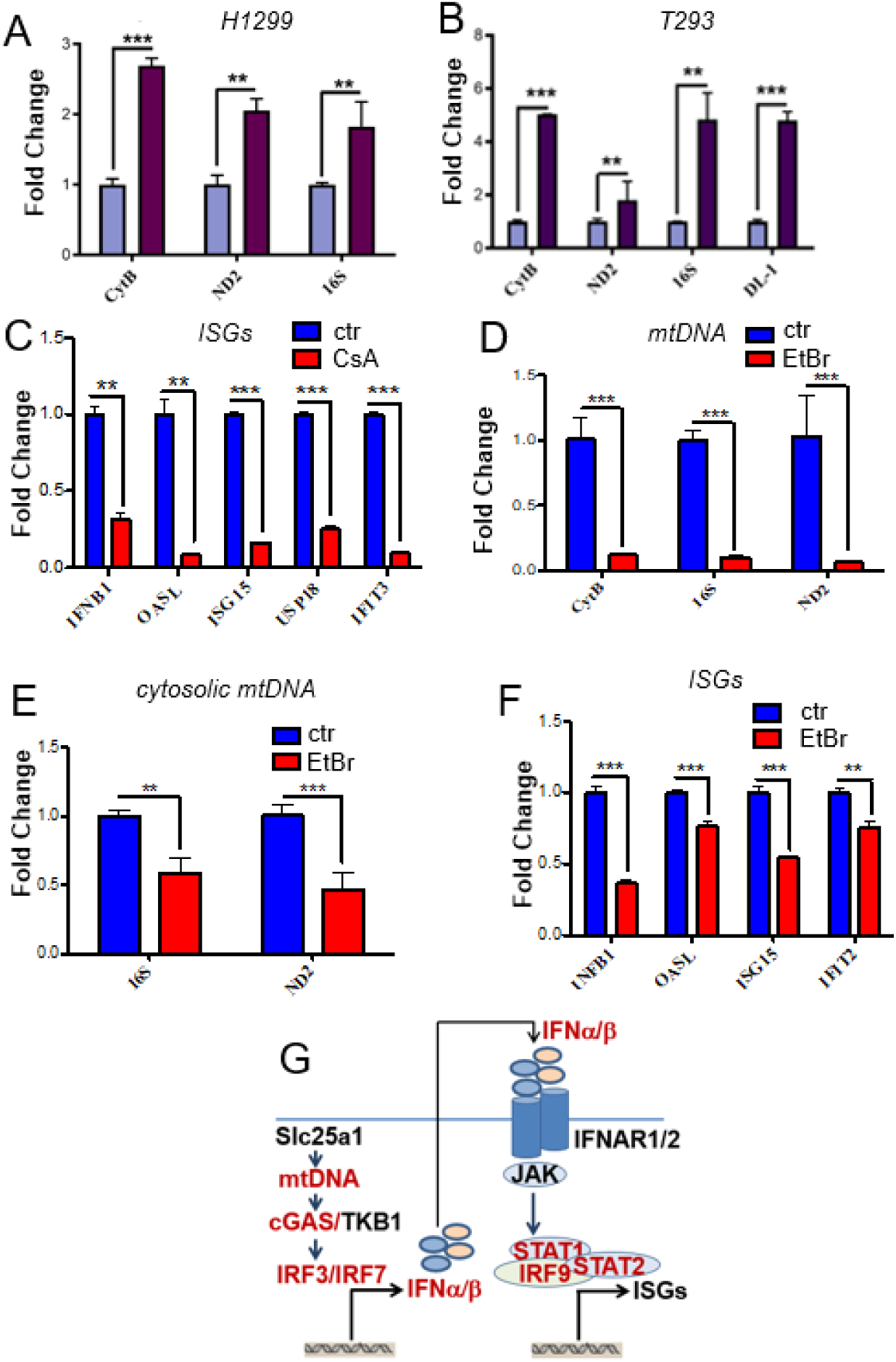
Cytosolic mtDNA accumulation drives the IFN-I response. **A-B.** Levels of mtDNA in the cytosolic fractions of SLC25A1^high^ H1299 (B) and T293 (C) cells (red, n=3) relative to control cells (blue, n=3). **C.** Fold change in gene expression of the indicated ISGs in SLC25A1^high^ H1299 spheres treated with CsA (red, n=3) relative to control untreated spheres (blue, n=3). **D.** Ratio of mtDNA to genomic DNA *(GAPD H*) measured by qRT-PCR on total extract of SLC25A1^high^ H1299 cells treated with EtBr (red, n=3) relative to control untreated cells (blue, n=3). **E.** Levels of mtDNA (16S and ND2) in the cytosolic fraction of SLC25A1^high^ H1299 cells treated with EtBr (red, n=3) relative to control untreated cells (blue, n=3). **F.** Fold change in gene expression of the indicated ISGs in SLC25A1^high^ H1299 spheres treated with EtBr (red, n=3) relative to untreated control spheres (blue, n=3). **G.** Schematic representation of the SLC25A1-IFN pathway and of the components that we found regulated (indicated in red). The error bars represent standard deviations, asterisks refer to *\*p* ≤ 0.05; *\*\*p* ≤ 0.01*; ***p* ≤ 0.001 by unpaired *T*-test.

mtDNA can exit the mitochondria *via* the Mitochondria Permeability Transition Pore (MPTP). Therefore, we inhibited the opening of the MPTP with Cyclosporine A (CsA), which suppresses pore opening by binding to cyclophilin D. We observed a significant reduction in the ISGs expression levels in CsA treated spheres relative to untreated cells (Fig. 3C), indicating a role for MPTP opening in the induction of the IFN-I response. To further confirm the involvement of mtDNA, we next studied the effect of mtDNA depletion on the induction of the ISGs by employing well-established protocols where mtDNA depletion was achieved by chronically treating cells with a very low concentration of Ethidium Bromide (EtBr) [e.g., 40]. At low concentrations, EtBr intercalates into mtDNA preventing replication, while having no observable effect on genomic DNA. mtDNA depletion was validated by quantifying the ratio of mtDNA to nuclear genomic DNA in total cell extracts using qRT-PCR (Fig. 3D). First, the assessment of the levels of mtDNA in the cytosolic fractions of EtBr-treated *versus* untreated cells demonstrated a marked decrease in cytosolic mtDNA accumulation in the treated cells, consistent with the observed total mtDNA depletion and as expected (Fig. 3E). Second, a clear reduction in ISG expression levels was observed in EtBr-treated cells, correlating with the depletion of total mtDNA and the diminished cytosolic accumulation of mtDNA (Fig. 3F). To exclude the possibility that EtBr treatment affects the expression of nuclear-encoded genes, we evaluated the expression of the housekeeping genes *Small Nuclear Ribonucleoprotein D3* (*SNRPD3*) and *TATA-Box Binding Protein* (*TBP*), and detected no significant differences between EtBr-treated and untreated cells (data not shown).

To validate that the cGAS/STAT1 pathway is induced by SLC25A1 downstream of cytosolic mtDNA release, we conducted shRNA-mediated knockdown of either cGAS or STAT1 and subsequently evaluated the ISG mRNA levels. The knockdown of either cGAS or STAT1 resulted in a marked reduction of ISG expression compared to cells transduced with control shRNA (Supplementary Fig. 2 B,C).

Together, these findings demonstrate that SLC25A1 promotes the release of mitochondrial DNA into the cytosol, triggering the cGAS-STAT1 pathway and establishing a virus-mimicry state that amplifies type I interferon signaling. This mechanism identifies SLC25A1 as a novel regulator of mitochondrial-driven immune signaling.

### SLC25A1 induces PD-L1 expression in a cell intrinsic and cell extrinsic fashion

In tumor cells, PD-L1 expression can be induced and amplified through autocrine and paracrine signaling loops involving interferons, cytokines and growth factors released by tumor cells within the tumor microenvironment [41–43]. Given the newly identified role of SLC25A1 in mediating immuno-regulatory pathways and IFN-I activation, we sought to determine whether SLC25A1 also influences PD-L1 expression levels in an autocrine and/or paracrine manner (depicted in Fig. 4A). We found that PD-L1 protein was markedly up-regulated in both SLC25A1^high^ H1299 and B16F0 cells (Fig. 4B and not shown). To rule out differences in the genetic background, we next assessed whether genetic or pharmacological inhibition of SLC25A1 influences PD-L1 expression in different cell lines. Treatment with the SLC25A1 inhibitor CTPI-2 significantly reduced PD-L1 protein levels, and the knockdown of SLC25A1 using a previously validated shRNA construct produced a similar decrease in the HCC827 lung cancer cell line, which naturally exhibits high endogenous PD-L1 expression. (Fig. 4C,D). Furthermore, the inhibitory effect of CTPI-2 on PD-L1 expression was also evaluated in the H1975 and HCC1937 cell lines, as well as in a patient-derived tumor cell line (T5) yielding similar results (Fig. 4E) and supporting the conclusion that, in certain cell types, PD-L1 expression is regulated by SLC25A1 activity. Given the fundamental role played by cell extrinsic signals, in a complementary experimental setting, we examined the effects of the culture supernatants on recipient cells. We found that PD-L1 expression was increased in cells exposed to the supernatants derived from SLC25A1-expressing spheres (Fig. 4F), demonstrating a critical role of this protein in regulation of PD-L1 expression.

**Figure 4.**
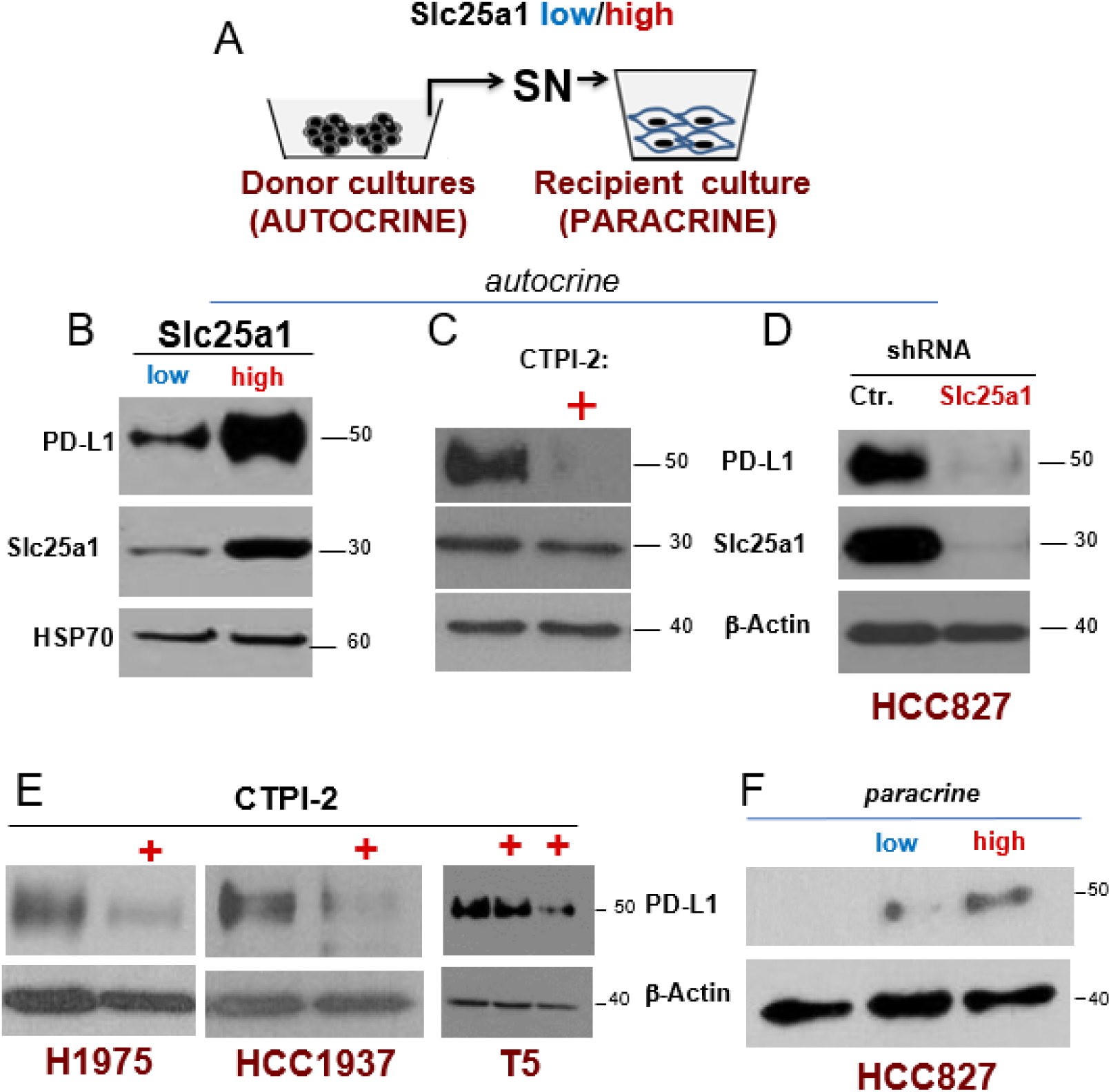
SLC25A1 induces PD-L1 expression in a cell intrinsic and extrinsic fashion. **A.** Schematic representation of the experiments. **B-D.** PD-L1 and SLC25A1 protein levels in control and SLC25A1^high^ H1299 spheres (B), or in spheres treated with CTPI-2 (C), or transduced with the SLC25A1 shRNA (D). **E.** PD- L1 and SLC25A1 protein levels in the indicated tumor (H1975 and HCC1937) and patient-derived (T5) NSLCs untreated or treated with CTPI-2. **F.** PD-L1 and SLC25A1 protein levels in HCC827 cells treated with control media, or with conditioned culture supernatant derived from SLC25A1^low^ or SLC25A1^high^ spheres.

### The SLC25A1-mediated induction of PD-L1 at least in part relies upon the ubiquitin adapter Keap1

Under normal physiological conditions, IFN-I up-regulates PD-L1 expression to promote immune homeostasis and prevent excessive inflammation and tissue damage. However, tumor cells exploit this pathway to promote immune evasion [41–43]. In keeping with the previous finding that SLC25A1 induces IFN-I signaling, we next explored whether the PD-L1 induction is dependent upon the IFN-I activation and the virus mimicry state triggered by SLC25A1. Contrary to this idea, mtDNA depletion led to PD-L1 upregulation, likely due to impaired mitochondrial respiration, which increases lactic acid production, known to increase PD-L1 levels (Fig. 5A) [44]. Similarly, the knock-down of the IFNAR1 receptor, cGAS or STAT1 only modestly reduced PD-L1 mRNA and protein levels in SLC25A1^high^ cells (Fig. 5B-D). Collectively, the data suggested the existence of additional molecular mechanisms by which SLC25A1 influences PD-L1 expression.

**Figure 5.**
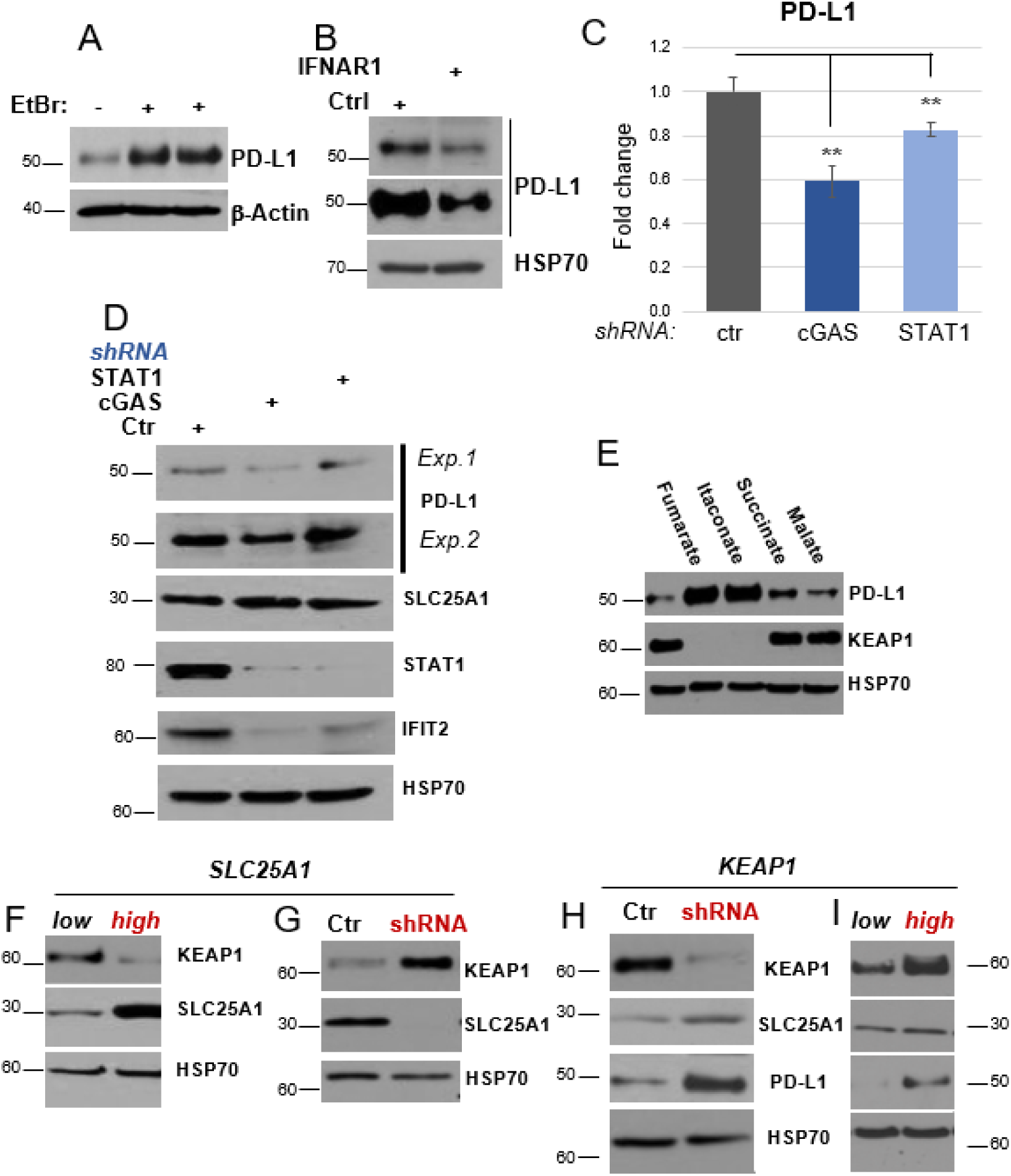
Induction of PD-L1 by fumarate and Keap1. **A.** PD-L1 protein levels in control or EtBr-treated H1299 spheres used at two different concentrations (indicated by +). **B.** PD-L1 protein levels in SLC25A1^high^ H1299 spheres infected with control lentivirus (pLKO) or with pLKO lentivirus harboring the shRNA for IFNAR1. **C-D**. Effects of the knock-down of cGAS or STAT1 on PD-L1 mRNA (C) or protein (D) levels in SLC25A1^high^ H1299 spheres. In D, two different exposures of the PD-L1 blot are shown (Exp. 1 and 2). **E**. Treatment of naïve H1299 with fumarate (DMF), itaconate, succinate and malate for 16 hours and immuno-blot with the indicated proteins. **F-G.** Keap1, PD-L1 and SLC25A1 protein levels in SLC25A1^high^ H1299 (F) or harboring the SLC25A1 shRNA (G). **H-I**. Keap1, PD-L1 and SLC25A1 protein levels in H1299 in spheres harboring control or the Keap1 shRNA (H), or, *viceversa*, over-expressing Keap1 (I, high), versus control cells.

We and others have previously shown that SLC25A1 promotes the tricarboxylic (TCA) cycle, enhancing the availability of several TCA cycle intermediates, including citrate, fumarate, malate, succinate, as well as of itaconate, an immunoregulatory TCA cycle metabolite derived from cis-aconitate. [34]. Given the emerging role of metabolic products and TCA cycle intermediates as immunoregulatory signaling metabolites capable of modulating inflammatory, hypoxic, and immune pathways linked to PD-L1 regulation, we investigated whether these metabolites contribute to the regulation of PD-L1 expression. This approach led to the discovery that treatment with either dimethyl fumarate (DMF), a cell-permeable methyl ester that releases fumarate as well as itaconate or, to a lesser extent, succinate, robustly upregulate PD-L1 protein levels (Fig. 5E), while citrate had no effect (not shown).

Fumarate and itaconate promote non-enzymatic post-translational modifications of cysteine residues *via* succination, resulting in the activation or inhibition of downstream targets. One of the best-characterized targets of this reaction is Keap1 (Kelch-like ECH-associated protein 1), the substrate adaptor of the BCR (BTB-Cul3-RBX1) E3 ligase complex. Succination of Keap1 by fumarate or itaconate leads to its functional inactivation and degradation. Consistent with this mechanism, DMF treatment markedly reduced Keap1 protein levels, coinciding with PD-L1 upregulation (Fig. 5E). In addition, SLC25A1 over-expression decreased Keap1 levels (Fig. 5F). Conversely, SLC25A1 knockdown resulted in Keap1 upregulation (Fig. 5G). Finally, the Keap1 knockdown enhanced PD-L1 protein expression (Fig. 5H), whereas Keap1 over-expression suppressed it (Fig.5I).

Thus, SLC25A1 enhances PD-L1 at least in part *via* downregulation of Keap1 by enhancing the availability of TCA cycle intermediates, particularly fumarate.

### SLC25A1 induces the IFN-I response and PD-L1 expression *in vivo* and sensitizes to anti-PD-L1 blockade

Building upon the results from our previous *in vitro* studies, we next investigated whether SLC25A1 influences the response to PD-L1 blockade. To accomplish this, we employed a syngeneic mouse model specifically the murine melanoma B16F0 cell line, a well-characterized and widely used model for studying immune checkpoint inhibitor (ICI) responses. Given the low basal, nearly undetectable expression of SLC25A1 in B16F0 cells (Fig.1C), we employed naïve B16F0 cells, as well as the stable derivative cell line that restores SLC25A1 protein levels (Fig. 1C). Control and SLC25A1^high^ cells were subcutaneously implanted into C57BL/6 mice, followed by treatment with either PD-L1-blocking antibodies or isotype control matched IgG, administered at 200 µg per mouse every three days for four doses once tumors became palpable. As amply reported in literature, parental B16F0 tumors exhibited minimal, if any, responsiveness to PD-L1/PD-1 blockade (Fig. 6 A,B). Consistent with the previously described role of SLC25A1 in promoting tumor growth, B16F0 cells stably expressing SLC25A1 had accelerated tumor growth rate compared to naïve cells. However, treatment with the PD-L1 antibody significantly suppressed the growth of SLC25A1^high^ tumors and reduced the incidence of progressing tumors in this group (Fig. 6 A).

**Figure 6.**
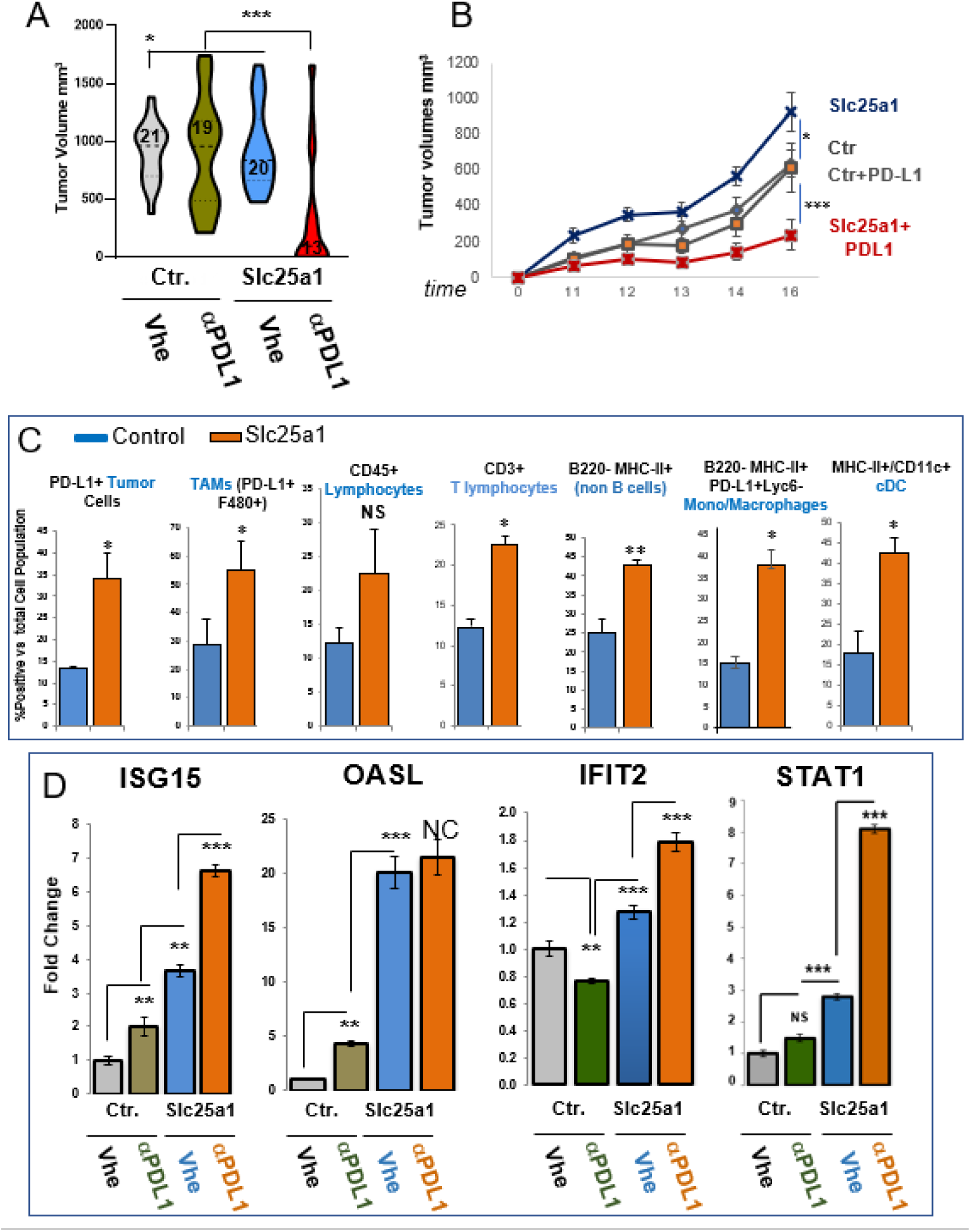
SLC25A1 influences the response to anti-PD-L1 antibody therapy. **A.** Tumor volumes for the indicated treatment groups from two independent experiments. **B.** Tumor growth curves of control and SLC25A1 over-expressing B16F0 tumors treated with either PD-L1 (αPD-L1) or isotype control antibodies. **C**. The abundance of the indicated cell populations were analyzed by FACS. The error bars represent the standard deviation, asterisks refer to *\*p* ≤ 0.05; *\*\*p* ≤ 0.01*; ** *p* ≤ 0.001 by unpaired *T*-test. **D.** q-RT-PCR of the indicated IFN-I markers.

Tumors from both groups were subsequently harvested for flow cytometry (FACS) analysis to characterize the immune infiltrate and assess PD-L1 expression and other markers of tumor immunogenicity, as described (Supplementary Fig.S3 and ref.45). Tumors derived from cells expressing SLC25A1 induced high levels of PD-L1 in both tumor cells and tumor-associated macrophages (TAMs) (Fig. 6C). This result is consistent with our previous *in vitro* findings showing autocrine and paracrine activities of the SLC25A1-mediated signal loop. Moreover, SLC25A1-expressing tumors exhibited increased infiltration of CD45+ leukocytes of CD3+ T cells, as well as enhanced expression of Histocompatibility Complexes positive monocytes and dendridic cells (Fig.6C), indicative of an inflammatory enriched microenvironment.

Considering the identified SLC25A1-dependent induction of IFN-I in our *in vitro* models, we next assessed the expression of representative sets of ISGs in these tumors. As shown in Fig. 6 D, a robust activation of IFN-I downstream targets, OASL, ISG15, IFIT2 and STAT1 was observed in SLC25A1^high^ tumors which was enhanced by the PD-L1 antibody.

Importantly, Kaplan–Meier survival analysis using the KMplotter database demonstrated that high PD-L1 expression was significantly associated with improved overall survival in patients treated with anti–PD-L1 immune checkpoint blockade (atezolizumab or durvalumab) (Fig.7A). Patients with high SLC25A1 expression exhibited a markedly prolonged survival probability compared with patients with low SLC25A1 expression (HR = 0.44, 95% CI: 0.32–0.60; log-rank *P* = 6.5 × 10⁻⁸), indicating an approximately 56% reduction in the risk of death in the high-expression group. These findings support the notion that elevated SLC25A1 expression may identify tumors more responsive to PD-L1–targeted immunotherapy and are consistent with the concept that SLC25A1 levels can serve as a predictive biomarker of response to checkpoint blockade.

**Figure 7.**
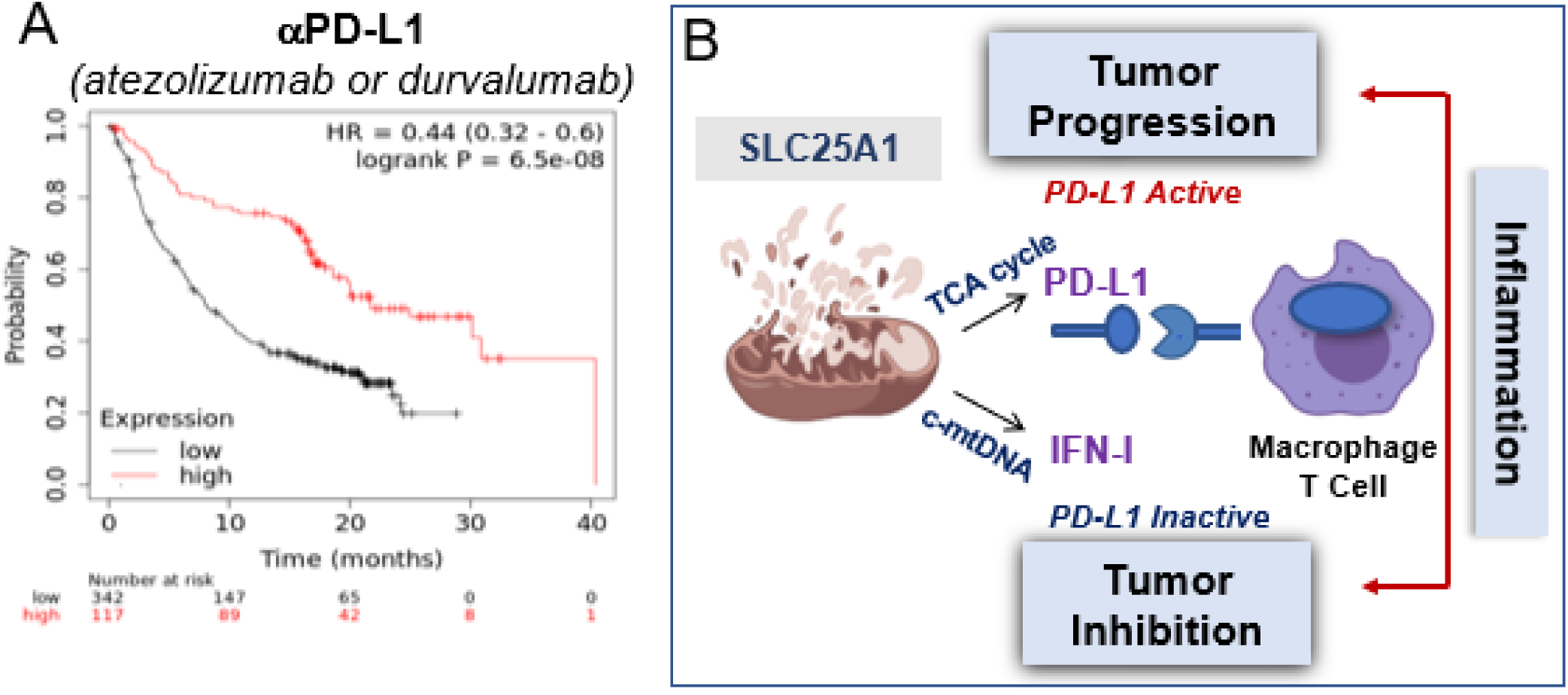
SLC25A1-Dependent Regulation of PD-L1 Activity and Association with Response to Anti–PD-L1 Immunotherapy. **A**. Kaplan–Meier overall survival curves of patients receiving anti–PD-L1 immune checkpoint therapy (atezolizumab or durvalumab) stratified according to PD-L1 expression levels using the KMplotter database. High PD-L1 expression was associated with significantly improved survival compared with low PD-L1 expression (HR = 0.44, 95% CI: 0.32–0.60; log-rank *P* = 6.5 × 10⁻⁸). No significant association was observed in patients treated with anti–PD-1 therapies. **B.** Proposed model of the tumor-initiated immuno-regulatory program driven by SLC25A1 and by the mitochondria that through autocrine and paracrine activities converge on regulation of PD-L1 and on the induction of a pro-inflammatory response (see also text).

## DISCUSSION

We describe a novel tumor-initiated immuno-regulatory program driven by SLC25A1 and by the mitochondria that through autocrine and paracrine activities converge on regulation of PD-L1 and on the induction of the IFN-I response (depicted in Fig. 7B). We first identified a novel molecular mechanism by which SLC25A1 induces a viral mimicry state and found that this induction is sustained by mtDNA, which in turn triggers activation of the IFN-I pathway. Second, SLC25A1 influences PD-L1 levels and, by investigating the molecular mechanisms underlying this cross-talk, we identified a previously unrecognized, novel mechanism regulating the expression of this protein, involving the activity of Keap1. Third, by employing syngeneic mouse models of tumor cells expressing high or low levels of SLC25A1, we show that melanoma xenografts with elevated SLC25A1 levels display a markedly improved response to PD-L1 blockade. Importantly, this enhanced response is associated with robust activation of ISGs. Therefore our data are consistent with the hypothesis that SLC25A1 creates a tumor environment that is primed for tumor killing but acquires “immune privilege” *via* up-regulation of PD-L1 and thus, these tumors are uniquely sensitive to PD-L1 inhibition. Mechanistically, we attribute this favorable outcome to pro-inflammatory activities supported by SLC25A1 that are “unleashed” once the inhibitory block imposed by PD-L1 is removed. In keeping with the pro-oncogenic roles played by inflammation, it is likely that this inflammatory response favors SLC25A1-mediated oncogenic program

Recent pan-cancer and large scale analyses have further highlighted the clinical significance of SLC25A1, corroborating the idea that its expression is broadly elevated across multiple tumor types and correlates with poor patient prognosis [47]. A recent study [47] further demonstrated that its high expression correlates with higher tumor grade, and more advanced disease stage. Elevated SLC25A1 levels were also associated with variable immune cell infiltration, immune checkpoint expression, and increased tumor mutational burden, further suggesting that SLC25A1 contributes to shaping the tumor microenvironment, a possibility that had not been explored directly up to this point.

As discussed previously, the effects of IFN-I in the response to immunotherapy are complex and at times conflicting. Indeed, chronic IFN-I induction has also been associated with resistance to ICIs in some instances. Based on our findings, we propose that a quantitative, dose-dependent effect of the IFN-I gene program may be required to mediate tumor killing, and we posit that SLC25A1, and the mitochondrial metabolism, acts by amplifying the magnitude of this response. Considering the pro-survival activities exerted by a subset of ISGs, it is also possible that the outcome of SLC25A1 activity and of ICIs is qualitatively influenced by the composition of the ISG gene program in a more granular fashion. Dissecting the composition of this ISG response remains an important area of ongoing investigation, and understanding which subsets of ISGs contribute to vulnerability *versus* resistance could allow for more precise therapeutic targeting.

Given that one major activity of SLC25A1 is the promotion of mitochondrial respiration [34], our data also raise the important question of whether drugs that stimulate mitochondrial OXPHOS might exert similar effects. Two such agents currently under investigation are dichloroacetate and fumarate, both of which are particularly attractive because they are already being evaluated in clinical trials. In addition, because tumor cells frequently exploit the PD-L1/PD1 axis and upregulate PD-L1 to evade anti-tumor immunity, there is considerable interest in understanding the mechanisms controlling PD-L1 stability. Indeed, elevated PD-L1 expression levels are often predictive of favorable clinical responses to ICIs; however, it is also documented that induction of PD-L1 degradation prevents immune escape and promotes recruitment of tumoricidal immune cells, thereby enhancing tumor killing in response to radio- or chemotherapy. In this respect, the identification of a novel pathway regulating PD-L1 expression levels involving both Keap1 and SLC25A1, provides an opportunity to rationally exploit SLC25A1-targeting agents to promote anti-tumor immune responses during conventional cancer therapies. We have shown here that CTPI-2 significantly down-regulates PD-L1 expression in several tumor cell lines and we and others have also demonstrated that in immuno-compromised mice, inhibition of the tumor-intrinsic activities of SLC25A1 using CTPI-2 synergizes with platinum therapy, radiotherapy, and EGFR-targeting agents. In light of the activities of SLC25A1 on the PD-L1 axis, these results establish a rationale for inhibiting SLC25A1 in combination with canonical anti-tumor therapies to prevent adaptive PD-L1 upregulation following chemo- or radiotherapy. Therefore, overall, our data suggest that there may be therapeutic benefit in targeting SLC25A1 activity either positively, when anti-PD-L1 represents the primary treatment, or negatively, when chemotherapy or targeted therapies are employed.

## MATERIALS AND METHODS

### Cells, reagents and antibodies

The H1299, B16F0, and H1937 cell lines were obtained from ATCC or from the tissue culture core facility at Lombardi comprehensive cancer center (LCCC) at Georgetown University. The 293T cell line was a kind gift from Dr. Chunling Yi. The HCC827 and H1975 cell line were kind gifts from Dr. Giuseppe Giaccone. The patient-derived NSCLC cell lines were described previously [34]. Cells were grown in Dullbecco’s modified Eagle’s medium (DMEM, 25 mM glucose, with glutamine and pyruvate from Invitrogen) and supplemented with 10% fetal bovine serum and 1% of 5000 units/ml of penicillin-streptomycin (pen-strep), (Gibco). The reagents used in this study were: the CTPI-2 is listed in Pubchem (CAS#: 68003-38-3) and it was purchased from Enamine Ltd, the Cyclosporine A (CsA) was purchased from Cell Signaling (9973), the Ethidium bromide was purchased from Alfa Aesar (J62282), the Dimethyl Fumarate was purchased from Sigma (242926), the Dimethyl Succinate was purchased from Sigma (W239607), the Dimethyl Malate was purchased from Sigma (374318), the MG132 was purchased from Sigma (474790), and the Chloroquine was purchased from Mpbio (193919). The vector-expressing human SLC25A1 was purchased from Origene (SC120727), and the vector-expressing Flag-PD-L1 was purchased from Sino Biological Inc. (HG10084-NF). The shRNA vectors used in this study were purchased from Sigma: SLC25A1 (TRCN0000232825), cGAS (TRCN0000149984), STAT1 (TRCN0000280021), IFNAR (TRCN0000059013) and Keap1 (TRCN0000154657). The anti-SLC25A1 antibody used in immuno-blot was either from Santa Cruz Biotech, (#SC-86392) or from Proteintech (15235-1-AP). Additional antibodies were as follows: PD-L1 (GeneTex, GTX104763), Keap1 (Proteintech, 10503-2-AP), OPA1 (Novus Biological, NB110-55290), cGAS (Cell Signaling, 15102), STAT1 (Cell Signaling, 14994), IFIT2 (Proteintech, 12604-1-AP), β-actin (Santa Cruz, sc-47778), and HSP-70 (Santa Cruz, sc-7298). Additional antibodies used for flow cytometry were: anti-mouse-CD45 (clone 30-F11, #564590), NK-1.1 (clone PK136, #562864), B220 (clone RA3-6B2, #563103), CD3e (clone 145-2C11, #564661), PD-L1 (clone MIH5, #564715), I-A/I-E (clone M5/114.15.2, #563415), CD11b (clone M1/70, #564443), CD11c (clone N418, #565452), all from BD Biosciences, and F4/80 (BM8, #123114), Gr-1 (clone RB6-8C5, #108428), and Ly6C (clone HK1.4, #128030) from BioLegend.

### Growth of monolayer and sphere cultures

Tumor cell lines were grown as monolayers in complete Dulbecco’s modified Eagle’s media (DMEM, Gibco; supplemented with 10% fetal bovine serum and 1% of 5000 units/ml of pen–strep, Gibco). To generate spheroids, cells were grown in falcon bacteriological Petri dishes coated with 2% poly(2-hydroxyethyl methacrylate) dissolved in 100% ethanol. The growth medium was DMEM/F12, Gibco; supplemented with 20 ng/ml of epidermal growth factor (EGF), 5 ng/ml of fibroblast growth factor, 0.375% 100× N2 supplements (Gibco) and 1% pen–strep. Monolayer cells were dissociated using 0.25% Trypsin-EDTA (Gibco), whereas spheroid cultures were dissociated using StemPro Accutase (Gibco).

### Inhibition of SLC25A1 activity

To pharmacologically inhibit SLC25A1 activity, stock solution of CTPI-2 was prepared by dissolving it in DMSO. CTPI-2 was added to the culture media of cells at a working concentration of 50 µM or 100 µM and incubation was carried out for 24 or 48 hours.

### Gene expression profiling of SLC25A1 over-expressing or control H1299 cells grown in monolayers or spheres conditions

Complete poly-A mRNA was purified from cell line samples using. Labeled cRNA was synthesized from cDNA per manufacturer instructions (Agilent). RNA was hybridized to an Agilent (Santa Clara, USA) SurePrint G3 Human Gene Expression v3 8×60K Microarray for quantification. Arrays were read using a SureScan microarray scanner. Per-spot intensity values were averaged by gene, quantile normalized, and background corrected using the R software package *limma.* Manhattan distances between individual samples and individual genes were used to create hierarchical clustering (Ward) per-samples and per-genes. Gene set enrichment was conducted using the R package *fgsea*.

### Immunoblot

Attached cells were harvested using cell scraper, spheres were harvested by centrifugation, and frozen tumor tissue samples were ground by mortar and pestle in liquid nitrogen. Cell lysis was performed using RIPA buffer supplemented with cOmplete Mini Protease Inhibitor Tablets (Roche, 11836153001). Protein quantification was done using Coomassie (Bradford) Protein Assay Kit (Pierce), equal amount of protein lysate was loaded and separated by Novex^™^ 4-20% Tris-Glycine Mini Gel (Invitrogen) then transferred to a PVDF membrane. The membrane was blocked in blocking buffer with 10% horse serum to prevent non-specific binding, followed by incubation with primary antibodies. Appropriate horseradish peroxidase–conjugated secondary antibody (Invitrogen) was applied after incubation with primary antibody, SuperSignal^™^ West Pico PLUS Chemiluminescent Substrate (Thermo Scientific) was used for protein detection. β-actin or HSP-70 was used as loading control.

### Quantitative real-time PCR

Total RNA was isolated using Trizol® (Thermo Scientific), and 5 µg of total RNA was treated with DNase I (Thermo Scientific), in the presence of SUPERase.In^TM^ RNase Inhibitor (Thermo Scientific) for 30 min at 37 °C; 5 mM EDTA was then added and the DNase I heat inactivated at 75 °C for 10 min. After adding 5 mM MgCl_2_, complementary DNA was generated using Superscript IV (Thermo Scientific) and random hexamers following the manufacturer’s instructions. Real-time PCR was carried out using PowerUp™ SYBR® Green Master Mix (Thermo Scientific), using the following reference genes for human samples: *GAPDH*, *SNRPD3* or *TBP* whereas for murine samples: actin, beta (*Actb*), Peptidyl-prolyl cis-trans isomerase A (*Ppia*) or *Tbp*. The gene expression was normalized to the average of two reference genes and the relative gene expression fold changes calculated using the ΔΔCT method. Dissociation curves were analyzed and showed single amplification products to confirm the specificity of each primer pair, and RT samples were run to verify no genomic DNA contamination was present.

### Sorting of CD133^+^ and CD44^+^/CD24^-^ CSCs

SLC25A1 over-expressing H1299 cells were grown in spheroid culture condition for three days to enrich for CSCs. Spheres were harvested by centrifugation, disassociated, washed with PBS and resuspended in DMEM. To isolate the CD133^+^ and CD133^-^ populations, disassociated spheres were stained with the APC anti-human CD133 antibody (BioLegend, #372806). To isolate the CD44^+^/CD24^-^ population, disassociated spheres were co-stained with the APC anti-mouse/human CD44 antibody (BioLegend, #103012) and the PE anti-human CD24 antibody (BioLegend, #311106). Cells were incubated with the antibodies at 37°C for 30 minutes, washed 3 times, and resuspended in DMEM. The sorting was carried out by the core facility of LCCC at Georgetown University. RNA was isolated form the sorted cell populations and the levels of the ISGs expression was assessed by qRT-PCR.

### Inhibition of MPTP opening

Cells grown in spheroid culture conditions were treated 15 µM CsA overnight. RNA was isolated and RT-PCR was performed to determine the expression levels the ISGs in control and treated spheres.

### Quantification of mtDNA levels in pure cytosolic fractions

Spheres were disassociated using StemPro Accutase (Gibco), resuspended in PBS to generate a single-cell suspension, and the cell count was determined. 5×10^6^ cells were suspended in 500 µl PBS. In a fresh microcentrifuge tube, 100 µl of the cell suspension were pelleted by centrifugation, resuspended in 500 µl of 50 µM NaOH, and placed on a heating block for 30 minutes to boil and solubilize DNA. Then, to normalize the pH in the generated cell extract, 50 µl of 1 M Tris-HCl pH 8 was added. This generated cell extract served as a normalization control for total mtDNA. The cells in the other remaining 400 µl of the cell suspension were pelleted by centrifugation and resuspended in 500 µl buffer made of 150 mM NaCl, 50 mM HEPES (pH 7.4), and 20 µg/ml digitonin (Sigma, D141). The cell suspension was incubated end over end at room temperature for 10 minutes to promote the selective permeabilization of plasma membrane. Then, intact cells were pelleted by centrifugation at 980 g for 3 minutes. The obtained cytosolic supernatants were transferred to a fresh microcentrifuge tube and the obtained pellet of intact cells was saved for immuno-blotting. To achieve a complete elimination of intact cells, the centrifugation step was repeated twice, transferring the cytosolic supernatant to a fresh microcentrifuge tube each time. A final centrifugation was done at 17000 g for 10 minutes to eliminate all the remaining cellular debris, generating a pure cytosolic fraction. Using QIAQuick Nucleotide Removal Kit (QIAGEN), DNA was isolated form the pure cytosolic fraction. Using primers designed to amplify regions of the *CYTB*, *ND2* and *16S* mitochondrial genes qRT-PCR was performed on DNA isolated from both whole cell extract and pure cytosolic fraction. The Ct values for the mitochondrial genes obtained from the cytosolic fraction were normalized the Ct values obtained from the whole cell extract to achieve optimal standardization and to account for variation in total mtDNA among the tested samples. The purity of the cytosolic fraction was confirmed by immuno-blotting, probing for a OPA1 as a mitochondrial protein marker.

### Depletion of mtDNA using EtBr

Cells were seeded in monolayer culture conditions, 150 ng/ml of EtBr was added to the culture media, and cells were allowed to grow for three days. Then, EtBr containing media was aspirated, cells were harvested by trypsinization and were seeded in spheroid culture conditions to from spheres. The generated spheres were pelleted by centrifugation, disassociated using StemPro Accutase (Gibco), resuspended in PBS to generate a single-cell suspension, and the cell count was determined. An equal number of cells form control and EtBr treated samples were resuspended in in 500 µl of 50 µM NaOH and placed on a heating block for 30 minutes to boil and solubilize DNA. To confirm the depletion of mtDNA, qRT-PCR was performed on DNA from total cell extract of both control and EtBr treated samples using primers designed to amplify the nuclear *GAPDH* gene as well as the mitochondrial *CYTB*, *ND2* and *16S* genes. The obtained Ct values for the mitochondrial genes were normalized to the Ct values obtained for the reference nuclear *GAPDH* gene. To confirm that EtBr had no effect on genomic DNA, the expression levels of the reference nuclear genes *SNRPD3* and *TBP* were determined by qRT-PCR.

### Conditioned culture supernatants experiments

Control and Slc25a21 over-expressing H1299 cells were seeded in spheroid culture conditions and allowed to condition the culture media for three days. Recipient HCC827 cells grown in monolayer culture conditions received unconditioned sphere culture media, conditioned culture supernatant derived from control spheres or conditioned culture supernatant derived from SLC25A1 over-expressing spheres. The HCC827 cells were allowed to grow for an overnight in each of the received culture media. Then, the recipient HCC827 cells were harvested, and immunoblotting was performed to determine the influence on PD-L1 protein levels.

### TCA metabolites treatments

Control H1299 cells were seeded in monolayer culture conditions and allowed to attach for an overnight. Then, cells were treated with 5 mM Dimethyl Malate, 5 mM Dimethyl Succinate or 5 mM Dimethyl Fumarate for an overnight. Control and treated cells were harvested, and immune-blotting was performed to determine the influence on PD-L1 protein levels.

### Syngeneic mouse model and ICIs treatment

Control and SLC25A1 over-expressing B16F0 cells were grown as spheres in DMEM/F12 media containing 2.5% KnockOut^TM^ Serum Replacement (Gibco), 1% glutamine, 1% 100x ITS liquid media supplement (Sigma, I3146), 1% Sodium Pyruvate, 0.1% Y-compound and 1% pen-strep for three days. Spheres were disassociated using StemPro Accutase (Gibco), resuspended in PBS to form single-cell suspensions and cell viability and count were determined. Five weeks old wild type C57BL/6J female mice were purchased from the Jackson Laboratory. Mice were subcutaneously injected with 5×10^5^ viable control or SLC25A1 over-expressing B16F0 cells in both right and left lower flanks. Once tumors became palpable, mice from the control and SLC25A1 overexpressing groups were randomized to receive anti-mouse PD-L1 antibody (BioXCell, BE0101) or rat IgG2b isotype control, anti-keyhole limpet hemocyanin (BioXCell, BE0090). Both the anti-mouse PD-L1 antibody and the isotype control were administered *via* intraperitoneal injections at concentration of 200 µg per mouse every three days for a total of four treatments. Tumors dimensions were routinely measured using a digital caliper, and tumor volumes were calculated using the following formula (length x width x hight x 0.523). All mice were sacrificed when the tumors reached the maximum allowed tumor volume, and all tumors were excised for subsequent analysis. All animal studies were approved by the Georgetown University Institutional Animal Care and Use Committee.

### Generation of single-cell suspension from solid tumor tissues

Extracted tumors were intensively chopped using sterile razor blades. The generated fragmented tumor tissues were resuspended in 10 ml DMEM media containing 1 mg/ml Collagenase D (Sigma, 11088866001) and passed multiple times through a 10 ml serological pipette. To obtain single-cell suspensions tumor tissue suspensions were placed in gentleMACS C Tubes (Miltenyi Biotec, 130-093-237) and the tumor tissues disassociation was carried out using the gentleMACS^TM^ Dissociator (Miltenyi Biotec, 130-095-937) using the program 37_m_TDK2. Post digestion, cell were filtered using 70 µm cell strainer and recovered by centrifugation, and subjected to FACS analysis.

### Multi-color flow cytometry analysis of immune cell infiltrates

Using the single-cell suspensions generated from control and SLC25A1 over-expressing B16F0 tumors, 2×10^6^ viable cells were aliquoted for staining. Then, cells were stained with a cocktail of antibodies for immune cells surface markers: anti-mouse PD-L1 (BD Pharmingen, 564715), anti-mouse F480 (BioLegend, 123114), anti-mouse CD45 (BD Horizon^TM^, 564590) and anti-mouse CD3e (BD Horizon^TM^, 564661). Cell were wash after 30 minutes staining incubation and analyzed using a FACS Symphony cytometer (BD Biosciences).

### Statistics

Statistical significance was assessed using both paired or unpaired, two-tailed Student t-test. Significant differences are indicated using the standard Michelin Guide scale (\**p* ≤ 0.05, significant; \*\**p* ≤ 0.01, highly significant; \*\*\**p* ≤ 0.001, extremely significant).

## Authors contribution

MLA designed most of the experiments and wrote the paper. RM designed several experiments, proof edited the manuscript for accuracy, performed the majority of the experiments, and organized the data for the Figures. MM performed the in vivo cell sorting experiments and MC provided important inputs for interpretation. AK and SP performed some of the experiments in the patient-derived cell lines and CH and CA contributed to the editing and development of the in vivo experiments.

## Acknowledgements and funding contributions

Studies on SLC25A1 in the MLA lab have been supported by R01CA193698, R21DE028670, R21CA256546 and R03TR004871. All these studies have been supported by the outstanding facility at GUMC and by the 2P30CA051008-30 CCSG grant. We are grateful to Dr. Harvey Fernandez, as well as to Anton Wellstein for many inputs on this project, to the Riegel/Wellstein and to Dr. Chunling Yi laboratories for sharing reagents and instrumentation. We also thank Dr. Giuseppe Giaccone, Dr. Pacini and Dr. Petrini, for the sharing of the patient-derived cell lines which were also described previously [34].

## Ethics statements

### Conflict of Interest

The authors declare that they have no known competing financial interests or personal relationships that could have appeared to influence the work reported in this paper.

### Ethical approval and consent to participate

All animal studies were conducted in compliance with ethical regulations according to protocol #2017-1192 approved by the Institutional Animal Care and Use Committee (IACUC) at Georgetown University. All the mice were housed at Georgetown University Division of Comparative Medicine, in a SPF vivarium which is maintained at a 12:12 h light:dark cycle, at 68–74°F and 30–70% humidity range, according to IACUC regulations. Mice were euthanized according to the IACUC guidelines. This study does not directly involve human subjects or human data that requires ethical approval. The collection and use of patient-derived samples utilized herein was approved as described previously [34].

### Data availability

All data will be made available upon request.

## Notes

### Competing Interest Statement

The authors have declared no competing interest.

### Summary of Updates

This a revision of the previous manuscript that has been updated with additional results, shown in Figure 6 and 7, and also includes some corrections from the previous version.

## REFERENCES

1. Sharma P, Hu-Lieskovan S, Wargo JA, Ribas A. Immune checkpoint therapy—current perspectives. Science 2023;374

2. Jiang Y, Chen M, Nie H, Yuan Y. PD-1 and PD-L1 in cancer immunotherapy: clinical implications and future considerations. Hum Vaccin Immunother 2019;15:1111–1122.

3. Wei SC, Duffy CR, Allison JP. Fundamental mechanisms of immune checkpoint blockade therapy. Cancer Discov 2018;8:1069–1086.

4. Petitprez F, Meylan M, de Reyniès A, Sautès-Fridman C, Fridman WH. The tumor microenvironment in the response to immune checkpoint blockade therapies. Front Immunol 2020;11:784.

5. Morad G, Helmink BA, Sharma P, Wargo JA. Hallmarks of response, resistance, and toxicity to immune checkpoint inhibitors. Cell 2021;184:5309–5327

6. Gibney GT, Weiner LM, Atkins MB. Predictive biomarkers for checkpoint inhibitor-based immunotherapy. Lancet Oncol 2016;17:e542–e551.

7. Yarchoan M, Albacker LA, Hopkins AC, et al. PD-L1 expression and tumor mutational burden are independent biomarkers of response to immune checkpoint inhibitors. J Clin Invest 2019;129:3374–3382.

8. Li H, Wang J, Huang L, et al. Biomarkers of response to PD-1 pathway blockade: evaluating PD-L1 expression as a predictor of ICI efficacy. Br J Cancer 2022;126:19–29.

9. Schaer DA, Beckmann RP, Dempsey JA, Huber L, Forest A, Amaladas N, et al. The CDK4/6 inhibitor abemaciclib induces a T cell-inflamed tumor microenvironment and enhances the efficacy of PD-L1 checkpoint blockade. Cell Reports 2018;22:2978–2994.

10. Zhang J, Bu X, Wang H, Zhu Y, Geng Y, Nihira NT, et al. Cyclin D-CDK4 kinase destabilizes PD-L1 via Cul3SPOP to control cancer immune surveillance. Nature 2018;553:91–95.

11. Gou Q, Dong C, Xu H, Khan B, Jin J, Liu Q, et al. PD-L1 degradation pathway and immunotherapy for cancer. Cell Death Dis 2020;11:955

12. Aricò E, Castiello L, Capone I, Gabriele L, Belardelli F. Type I interferons and cancer: an evolving story demanding novel clinical applications. Cancers (Basel) 2019;11:1943.

13. Zemek RM, De Jong E, Chin WL, Schuster IS, Fear VS, Casey TH, et al. Sensitizing the tumor microenvironment to immune checkpoint therapy. Front Immunol 2020;11:223.

14. Medrano RFV, Hunger A, Mendonça SA, Barbuto JAM, Strauss BE. Immunomodulatory and antitumor effects of type I interferons and their application in cancer therapy. Oncotarget 2017;8:71249–71284.

15. Minn AJ, Wherry EJ. Combination cancer therapies with immune checkpoint blockade: convergence on interferon signaling. Cell 2016;165:272–275.

16. Teijaro JR. Type I interferons in viral control and immune regulation. Curr Opin Virol 2016;16:31–40

17. Qadir AS, Ceppi P, Brockway S, Law C, Mu L, Khodadadi-Jamayran A, et al. CD95/Fas increases stemness in cancer cells by inducing a STAT1-dependent type I interferon response. Cell Rep 2017;18:2373–2386.

18. Erdal E, Haider S, Rehwinkel J, Harris AL, McHugh PJ. A prosurvival DNA damage-induced cytoplasmic interferon response is mediated by end resection factors and is limited by Trex1. Genes Dev 2017;31:353–369.

19. Weichselbaum RR, Ishwaran H, Yoon T, Nuyten DS, Baker SW, Khodarev N, et al. An interferon-related gene signature for DNA damage resistance is a predictive marker for chemotherapy and radiation for breast cancer. Proc Natl Acad Sci U S A 2008;105:18490–18495.

20. Benci JL, Xu B, Qiu Y, Wu TJ, Dada H, Twyman-Saint Victor C, et al. Tumor interferon signaling regulates a multigenic resistance program to immune checkpoint blockade. Cell 2016;167:1540–1554.e12.

21. Benci JL, Johnson LR, Choa R, Xu Y, Qiu J, Zhou Z, et al. Opposing functions of interferon coordinate adaptive and innate immune responses to cancer immune checkpoint blockade. Cell 2019;178:933–948.e14.

22. Guo J, Zhou J, Yuan Y, Huang Y, Yang J, Qiu Y, et al. Empowering therapeutic antibodies with IFN-α for cancer immunotherapy. PLoS One 2019;14:e0219829.

23. Romero D. Interferon enhances immune-checkpoint inhibition. Nat Rev Clin Oncol 2019;16:6.

24. Zhu Y, Tibensky I, Schmidt J, Ryschich E, Märten A. Interferon-α enhances antitumor effect of chemotherapy in an orthotopic mouse model for pancreatic adenocarcinoma. J Immunother 2008;31:599–606.

25. Harel M, Ortenberg R, Varanasi SK, Mangalhara KC, Mardamshina M, Markovits E, et al. Proteomics of melanoma response to immunotherapy reveals mitochondrial dependence. Cell 2019;179:236–250.e18.

26. Gonzalez-Cao M, Iduma P, Karachaliou N, Santarpia M, Blanco J, Rosell R. Activation of viral defense signaling in cancer. Ther Adv Med Oncol 2018;10:1758835918793108.

27. Deblois G, Smith HW, Tam IS, Gravel SP, Caron M, Savage P, et al. Epigenetic switch–induced viral mimicry evasion in chemotherapy-resistant breast cancer. Cancer Discov 2020;10:1312–1329.

28. Roulois D, Loo Yau H, Singhania R, Wang Y, Danesh A, Shen SY, et al. DNA-demethylating agents target colorectal cancer cells by inducing viral mimicry by endogenous transcripts. Cell 2015;162:961–973.

29. Formenti SC, Rudqvist NP, Golden E, Cooper B, Wennerberg E, Lhuillier C, et al. Radiotherapy induces responses of lung cancer to CTLA-4 blockade. Nat Med 2018;24:1845–1851.

30. Won JK, Bakhoum SF. The cytosolic DNA-sensing cGAS–STING pathway in cancer. Cancer Discov 2020;10:26–39.

31. Bakhoum SF, Ngo B, Laughney AM, Cavallo JA, Murphy CJ, Ly P, et al. Chromosomal instability drives metastasis through a cytosolic DNA response. Nature 2018;553:467–472.

32. Kasprzyk-Pawelec A, Tan M, Rahhal R, McIntosh A, Fernandez HR, Mosaoa RM, Jiang L, Pearson GW, Glasgow E, Vockley J, Albanese C, Avantaggiati ML. Inactivation of the SLC25A1 gene during embryogenesis induces a unique senescence program controlled by p53. Cell Death Differ 2025;32:818–836.

33. Mosaoa R, Kasprzyk-Pawelec A, Fernandez HR, Avantaggiati ML. The mitochondrial citrate carrier SLC25A1/CIC and the fundamental role of citrate in cancer, inflammation and beyond. Biomolecules 2021;11:141.

34. Fernandez HR, Gadre SM, Tan M, Graham GT, Mosaoa R, Ongkeko MS, et al. The mitochondrial citrate carrier, SLC25A1, drives stemness and therapy resistance in non-small cell lung cancer. Cell Death Differ 2018;25:1239–1258.

35. Catalina-Rodriguez O, Kolukula VK, Tomita Y, Preet A, Palmieri F, Wellstein A, et al. The mitochondrial citrate transporter, CIC, is essential for mitochondrial homeostasis. Oncotarget 2012;3:1220–1235.

36. Butow RA, Avadhani NG. Mitochondrial signaling: the retrograde response. Mol Cell 2004;14:1–15.

37. Liu Z, Butow RA. Mitochondrial retrograde signaling is a pathway of communication from mitochondria to the nucleus under normal and pathophysiological conditions. Annu Rev Genet 2006;40:159–185.

38. Bhat N, Fitamant J, Hauck L, et al. Recognition of cytosolic DNA by cGAS and other STING pathway sensors. Nat Rev Immunol 2014;14:76–88.

39. Kong LZ, Xue WJ, Wang ZH, et al. Understanding nucleic acid sensing and its therapeutic opportunities. Mol Cancer 2023;22:180.

40. Wu Z, Bogardus H, Safaee Modaress S, Esposito D, Caveloo H, Shadel GS. Mitochondrial DNA stress signalling protects the nuclear genome. Nat Commun 2019;10:5646.

41. Sun C, Mezzadra R, Schumacher TN. Regulation and function of the PD-L1 checkpoint. Immunity 2018;48:434–452.

42. Jiang X, Wang J, Deng X, Xiong F, Ge J, Xiang B, et al. Role of the tumor microenvironment in PD-L1/PD-1-mediated tumor immune escape. Mol Cancer 2019;18:10.

43. Cha JH, Chan LC, Li CW, Hsu JL, Hung MC. Mechanisms controlling PD-L1 expression in cancer. Mol Cell 2019;76:359–370.

44. Chen J, Chen Z, Chen M, Liu H, Luo Y, Xu L, et al. Tumor cell-derived lactate induces TAZ-dependent upregulation of PD-L1. Oncogene 2017;36:5829–5839.

45. Ajina R, Malchiodi ZX, Fitzgerald AA, Zuo A, Wang S, Moussa M, et al. Antitumor T-cell immunity contributes to pancreatic cancer immune resistance. Cancer Immunol Res 2021;9:386–400.

46. Nasrollahzadeh E, Rahbarizadeh F, Ahmadvand D, Hartmann T. Pro-tumorigenic functions of macrophages at the primary, invasive and metastatic tumor site. Cancer Immunol Immunother 2020;69:1673–1697.

47. You Y, Wu Y, Chen Y, Fang H, Xu Y, Dai Y, Zhang J, Wu S. A comprehensive analysis of SLC25A1 expression and its prognostic potential in cancers. Front Oncol. 2023;13:1224995.

